# *ZAXINONE SYNTHASE 2* regulates growth and arbuscular mycorrhizal symbiosis in rice

**DOI:** 10.1101/2022.07.21.501002

**Authors:** Abdugaffor Ablazov, Cristina Votta, Valentina Fiorilli, Jian You Wang, Fatimah Aljedaani, Muhammad Jamil, Aparna Balakrishna, Raffaella Balestrini, Kit Xi Liew, Chakravarthy Rajan, Lamis Berqdar, Ikram Blilou, Luisa Lanfranco, Salim Al-Babili

**Affiliations:** King Abdullah University of Science and Technology (KAUST), Biological and Environmental Sciences and Engineering Division, Center for Desert Agriculture (CDA), The BioActives Lab, Thuwal 23955- 15 6900, Saudi Arabia; Department of Life Sciences and Systems Biology, University of Torino, Viale Mattioli 25, Torino 10125, Italy; Plant Science Program, Biological and Environmental Science and Engineering Division, King Abdullah University of Science and Technology (KAUST), Saudi Arabia; Plant Cell and Developmental Biology, Biological and Environmental Sciences and Engineering (BESE), King Abdullah University of Science and Technology (KAUST), Thuwal 23955-6900, Saudi Arabia; Institute for Sustainable Plant Protection – CNR, Strada delle Cacce 75, 10135, Torino, Italy

**Keywords:** Apocarotenoid, Arbuscular Mycorrhizal fungi symbiosis, CAROTENOID CLEAVAGE DIOXYGENASES, Strigolactone, Zaxinone Synthase

## Abstract

Carotenoid cleavage, catalyzed by CAROTENOID CLEAVAGE DIOXYGENASES (CCDs), provides signaling molecules and precursors of plant hormones. Recently, we showed that zaxinone, a novel apocarotenoid metabolite formed by the CCD Zaxinone Synthase (ZAS), is a growth regulator required for normal rice growth and development. The rice genome encodes three *OsZAS* homologs, called here *OsZAS1b, OsZAS1c*, and *OsZAS2*, with unknown functions. Here, we investigated the enzymatic activity, expression pattern, and subcellular localization of OsZAS2, and generated and characterized loss-of-function CRISPR/Cas9-*Oszas2* mutants. We show that OsZAS2 formed zaxinone *in vitro*. OsZAS2 is a plastid-localized enzyme mainly expressed in the root cortex under phosphate starvation. Moreover, *OsZAS2* expression increased during mycorrhization, specifically in arbuscule-containing cells. *Oszas2* mutants contained lower zaxinone content in roots and exhibited reduced root and shoot biomass, less productive tiller, and higher strigolactone (SL) levels. Exogenous zaxinone application repressed SL biosynthesis and partially rescued the growth retardation of *Oszas2* mutant. Consistent with the *OsZAS2* expression pattern, *Oszas2* mutants displayed a lower frequency of AM colonization. In conclusion, *OsZAS2* encodes a further zaxinone-forming enzyme that determines rice growth and architecture and strigolactone content and is required for optimal mycorrhization.

## Introduction

Carotenoids are tetraterpene (C_40_) pigments consisting of long hydrocarbon chains with a conjugated double-bond system. In plants, carotenoids serve as a crucial component of photosynthesis, colorants, and antioxidants (Fraser and Bramley, 2004; Ballottari et al., 2014; Nisar et al., 2015; Hashimoto et al., 2016; Rodriguez et al., 2018; Zheng et al., 2020). In addition, the breakdown of carotenoids gives rise to a diverse group of metabolites called apocarotenoids, which includes pigments, scents, signaling molecules, growth regulators, and the precursors of the phytohormones strigolactone (SL) and abscisic acid (ABA) (Felemban et al., 2019; Moreno et al., 2021; Wang et al, 2021a; Liang et al., 2021; Zheng et al., 2021). ABA is the most-studied plant apocarotenoid hormone and a key player in plant response to abiotic and biotic stress (Peleg and Blumwald, 2011), regulation of seed maturation, dormancy, and shoot and root growth (Nambara and Marion-Poll, 2005; Moreno et al., 2021). SLs regulate a series of developmental processes. They are best known for inhibiting shoot branching/tillering, regulating root architecture, secondary growth, and senescence, and their contribution to biotic and abiotic stress responses (Gomez-Roldan et al., 2008; Umehara et al., 2008; Al-Babili and Bouwmeester, 2015; Waters et al., 2017; Jia et al., 2019; Van Ha et al., 2014; Decker et al., 2017). However, SLs were originally discovered as the host root-released germination stimulants for seeds of root parasitic weeds (Xie and Yoneyama, 2010). Later on, they were identified as the plant-released hyphal branching factor for arbuscular mycorrhizal (AM) fungi, which paves the way for establishing plant-AM symbiosis (Akiyama et al., 2005). AM fungi symbiotic association provides the host plant with minerals, mainly phosphorus (P) and nitrogen (N), and the AM fungi with carbohydrates and lipids (Wang et al., 2017). AM symbiosis is widely distributed and formed by most of the land plants, mirroring its importance for their growth and survival (Gutjahr and Parniske, 2013; Wang et al., 2017; Fiorilli et al., 2019).

Recent studies unraveled several new apocarotenoid signaling molecules, such as anchorene, iso-anchorene, β-cyclocitral, and zaxinone. Anchorene is a carotenoid-derived dialdehyde responsible for anchor root formation in *Arabidopsis thaliana* (Jia et al., 2019), while its structural isomer iso-anchorene inhibits Arabidopsis root growth (Jia et al., 2021). β-Cyclocitral regulates root growth and is a retrograde signaling molecule that mediates a single oxygen response and improves the high light tolerance by modulating the expression of oxidative stress-responsive genes (Ramel et al., 2012; Dickinson et al., 2019). Zaxinone is a regulatory metabolite, which is required for normal rice growth and development and negatively regulates SL biosynthesis (Wang et al., 2019; Wang et al., 2020). Multi-omics study revealed that zaxinone also modulates cytokinin homeostasis and that its growth-promoting effect is likely caused by increased sugar metabolism in rice roots (Wang *et. al*, 2021). However, exogenous application of zaxinone to Arabidopsis simultaneously increased both SL and ABA content (Ablazov et al., 2020), suggesting that it might act as a stress signal in Arabidopsis (Ablazov et al., 2020).

Apocarotenoid production in plants is mediated by carotenoid cleavage dioxygenases (CCDs), which cleave double bonds in carotenoid backbones and exhibit different substrate and cleavage site specificities. The diversity of CCDs gives rise to a wide spectrum of apocarotenoids with unique features and functions (Guiliano et al., 2003; Auldridge et al., 2006; Ahrazem et al., 2016). Based on phylogenetic analysis and enzymatic activity, plant CCDs build six subfamilies; NINE-*CIS*-EPOXY CAROTENOID DIOXYGENASES (NCEDs), CCD1, CCD4, CCD7, CCD8, and ZAXINONE SYNTHASE (ZAS) (Wang et al., 2019). NCEDs are involved in ABA biosynthesis and catalyze the cleavage of the C11, C12 (or C11’, C12’) double bond in *9-cis-epoxy* carotenoids to yield the ABA precursor xanthoxin (Schwartz et al., 1997; Jacqueline et al., 2000). In contrast to other CCD-types investigated so far, CCD1 enzymes are localized in the cytosol. Moreover, they are characterized by wide substrate and regio-specificity, cleaving many carotenoid and apocarotenoid substrates and producing dialdehyde products and volatiles that contribute to the flavor and aroma in many plants (Vogel et al., 2008; Ilg et al., 2009; Ilg et al., 2014). CCD4 enzymes cleave the C9-C10 or C9’-C10’ double bond in bicyclic carotenoids, giving rise to C_13_ volatiles and C_27_-apocarotenoids (Bruno et al., 2015; Bruno et al., 2016). CCD7 and CCD8 act sequentially on 9-*cis*-β-carotene to produce carlactone, the central intermediate of SL biosynthesis, via the intermediate 9-*cis*-β-apo-10’-carotenal formed by CCD7 along with the volatile β-ionone (Alder et al., 2012; Bruno et al., 2014). Carlactone is further modified by cytochrome P450s (711 clades), such as the *Arabidopsis* MORE AXILLARY GROWTH1 (MAX1) or the rice carlactone oxidase (CO), leading to the formation of canonical and non-canonical SLs (Abe at al., 2014; Zhang et al., 2014; Jia et al., 2018; Ito et al., 2022).

ZAS is a newly discovered member of the CCD family, which cleaves the apocarotenoid C27 apo-10-zeaxanthinal (C_27_) at the C13, C14 double bond, forming the C_18_-apocarotenoid zaxinone (Wang et al., 2019). Zaxinone is a growth-promoting metabolite required for normal rice growth and a negative regulator of SL biosynthesis and release (Wang et al., 2019; Wang et al., 2020). A rice loss-of-function *zas* mutant showed reduced root zaxinone level, retarded growth, i.e. lower root and shoot biomass, tiller number, and higher SL content (Wang et al., 2019). Confirming its biological function, the exogenous application of zaxinone restored several phenotypes of the *zas* mutant (Wang et al., 2019). Though all other CCD subfamilies are conserved, non-mycorrhizal species, such as *A. thaliana* and other members of the *Brassicales*, lack *ZAS* orthologues, indicating a role of *ZAS* in AM-symbiosis (Fiorilli et al., 2019; Wang et al., 2019). Indeed, the *zas* mutant displayed a lower level of AM colonization compared to the wild-type (Wang et al., 2019).

The rice genome contains three *OsZAS* homologs, previously called *OsZAS-L1* (renamed *OsZAS1b), OsZAS-L2* (renamed *OsZAS1c*), and *OsZAS-L3* (renamed *OsZAS2*) with unknown functions (Wang et al., 2019). In this study, we investigated the biological function of OsZAS2 by studying its enzymatic activity and generating and characterizing corresponding mutant and GUS reporter lines. Obtained data suggest that OsZAS2 is a non-redundant, root- and arbusculated cell-localized zaxinone-forming enzyme required for proper growth and normal SL homeostasis and mycorrhization level.

## Materials and Methods

### Plant material and phenotyping

Rice seedlings (*Oryza Sativa* L. cv Dongjin) were grown in a Biochamber under the following conditions: a 12 h photoperiod, 500-μmol photons m^-2^ s^-1^ and day/night temperature of 27/25 ° C. Briefly, rice seeds were surface sterilized in a 50% household bleach for 15 min and rinsed five times with distilled water. Then, sterilized seeds were germinated in the dark for 2 days in the magenta boxes containing 50 ml of 0.4% agarose half-strength Hoagland medium with pH 5.8 at 30 °C. The pre-germinated seeds were transferred to the biochamber and kept for 5 days.

For metabolites, gene expression, and *Striga* seed germination assay, one-week-old rice seedlings were transferred into 50 ml black falcon tubes filled with half-strength modified Hoagland nutrient solution with adjusted pH to 5.8. The nutrient solution consisted of 5.6 mM NH_4_NO_3_, 0.8 mM MgSO_4_.7H_2_O, 0.8 mM K_2_SO_4_, 0.18 mM FeSO_4_.7H_2_O, 0.18 mM Na_2_EDTA.2H_2_O, 1.6 mM CaCl_2_.2H_2_O, 0.8 mM KNO_3_, 0.023 mM H_3_BO_3_, 0.0045 mM MnCl_2_.4H_2_O, 0.0003 mM CuSO_4_.5H_2_O, 0.0015 mM ZnCl_2_, 0.0001 mM Na_2_MoO_4_.2H_2_O and 0.4 mM K_2_HPO_4_.2H_2_O. For normal conditions (+Pi), the one-week-old seedlings were grown in the Hoagland nutrient solutions (+Pi) for another two weeks. For phosphate starvation, the seedlings were grown for two weeks in lower phosphate (4 μM, K_2_HPO_4_·2H_2_O) nutrient solution. The nutrient solution was replaced every three days. For zaxinone treatment, three weeks-old seedlings were treated with 5 μM of zaxinone for 6 h: tissues were collected and immediately frozen into liquid N_2_.

For phenotyping, one-week-old *Oszas2* mutants seedlings were transferred into pots filled with soil and grown in growth chamber conditions as above. Tap water was supplied when needed. After 18 days, the seedling’s root was separated from the soil. Then, the seedlings were pictured with a digital camera, and root and shoot lengths were measured. To analyze the dry weight (DW) of root and shoot biomass, samples were kept for 3 days in a 65° C oven. For phenotyping of *Oszas2* mutants at the mature stage, one-week-old seedlings were transferred into a greenhouse and grown until the mature stage with a day/night temperature of 28/25 °C. One-time tap water and a one-time half-strength nutrient solution were supplied when necessary. After three months, yield-related traits were recorded.

For zaxinone rescue of *Oszas2* mutant, 7 days old seedlings were transferred into one-liter pots filled with soil and grown in Biochamber. Initially, for zaxinone treatment 200 ml of 10 μM of zaxione solutions (pH5.8) and for control same volume of tap water (0.01% Acetone) were added per pot. Every three days, 50 ml of 10 μM of zaxione solution and an equivalent amount of water were added to the treatment and control groups, respectively. After two weeks of treatment, the seedlings were phenotyped as above.

### Generation of transgenic lines

The CRISPR/Cas9 genome-editing technique was used to knock out *OsZAS2 (Os06g0162550*) in *Oryza sativa ssp. japonica* variety Dongjin. The gRNA sequences were designed using the CRISPR-PLANT webserver (www.genome.arizona.edu/crispr/). Two different gRNA sites targeted *OsZAS2* in exon regions; at exon 1 which encodes (5’-GGTCACTAGATGCATTCATC-’3) and exon 2 which encodes (5’-GCAGATCTGAAGAGACTGAT-’3). The gRNA spacers were fused to a tRNA sequence as described by Xie *et al*. (2015) (Supplemental Figure S5) and synthesized from GENEWIZ (South Plainfield, NJ, USA). Then, the sequence was cloned into pRGEB32 (Kanamycin) via 5’BsaI and 3’BsaI. The *pRGEB32-OsZAS2* construct was further introduced into the *Agrobacterium tumefaciens* strain *EHA105* competent cells *via* electroporation. To construct *the pOsZAS2::GUS* reporter plasmid, the 1.2 kb promoter region of *OsZAS2* was amplified with the Phusion enzyme from the genomic DNA of rice using promoter-specific primers (Supplemental Table S1). The PCR product ligated to pJet1.2 intermediate plasmid following the instruction of CloneJET PCR Cloning Kit (K1232, Thermo Scientific). The *pOsZAS2* sequence was amplified from the pJet1.2 plasmid with the specific primers (Supplemental Table S1) and cloned into the pENTR™/D-TOPO™ plasmid. Then, OsZAS2:pENTR™/D-TOPO™ were inserted into pMDC162 by Gateway cloning.

Rice transformation was conducted according to Hiei and Komari (2008). The mutations of transformed lines were analyzed by PCR using a Thermo Scientific™ Phire Plant Direct PCR Master Mix Kit. Gene-specific primers (Supplemental Table S1) were used for PCR amplification to detect the mutation sites. Then, PCR products were cleaned up using ExoSAP-IT™ PCR Product Cleanup Reagent and submitted to the Sanger sequencing core lab team, at KAUST. The Sanger sequencing data (abi file) were analyzed following the instruction of DSDecode (Degenerate Sequence Decode; http://skl.scau.edu.cn/dsdecode/). Three independent homozygote mutant lines were identified and grown until T3 generation.

### Metabolite quantification

For quantification, plant material was lyophilized with Freeze-dryer and ground with Geno Grinder 2010. D_3_-zaxinone (customized synthesis; Buchem B.V., Apeldoorn, The Netherlands) was used as an internal standard (IS). Zaxinone was extracted according to Wang *et al*. (2019). SLs extracted from the root tissue as described by Mi *et al*. (2018). The SLs extraction from the root exudate was performed according to Wang et al. (2022). In the final step, the dried extract was dissolved in 110 μL of acetonitrile: water (90:10, v:v) and filtered through a 0.22 μm filter for LC-MS/MS analysis. The samples were run on UHPLC-Triple-Stage Quadrupole Mass Spectrometer (TSQ-Altis™) with parameters as described in Wang et al. (2022).

### *Striga* seed germination bioassays

*Striga* seeds were pre-conditioned as described in Jamil et al. (2019). After 10 days, the pre-conditioned *Striga* seeds were treated with rice root extracts and exudates, at 50 μl per disc (n=3-6). Root extract prepared as the above SL extraction method and 5 μl of root extract diluted in 400 μl of MilliQ water before application. Root exudate collected as above SL root exudate collection protocol and 200 μl SL enriched solution diluted in 1800 μl of MilliQ water before application. The discs were also treated with water and GR24 (1μM) to include negative and positive control. The plates were again sealed with parafilm and incubated at 30 °C for 24 h. The discs were scanned in a microscope and germinated and non-germinated seeds were counted from these scanned images by using the software SeedQuant (Braguy et al., 2021) to calculate the percentage of germination.

### Real-time PCR analysis

Rice tissues were ground and homogenized in liquid nitrogen, and total RNA was isolated using a Direct-zol RNA Miniprep Plus Kit following the manufacturer’s instructions (ZYMO RESEARCH; USA). Briefly, a 1-μg RNA sample was reverse transcribed using an iScript cDNA Synthesis Kit (BIO-RAD Laboratories, Inc, 2000 Alfred Nobel Drive, Hercules, CA; USA). The qRT-PCR was performed using SYBR Green Master Mix (Applied Biosystems; www.lifetechnologies.com) in a CFX384 Touch™ Real-Time PCR Detection System (BIO-RAD Laboratories, Inc, 2000 Alfred Nobel Drive, Hercules, CA; USA). Primer-BLAST webserver (Ye et al., 2012) was used to design the gene-specific RT-qPCR primers (Supplemental Table S1) and Ubiquitin (OsUBQ) used as an internal control. The relative gene expression level was calculated according to 2^Δ^CT method.

### *In vitro* assays

The OsZAS2 cDNA was amplified using primers listed in Supplemental Table S1 and cloned into the pET-His6 MBP N10 TEV LIC cloning vector (2C-T vector; http://www.addgene.org/29706/) with MBP tags at the N-terminus. OsZAS2-MBP construct transformed into the BL21 Rosetta *E.coli* cells. A single colony of the transformed *E.coli* was cultured overnight and 0.5 mL of this culture inoculated into 50 mL liquid media and grown at 37°C to OD 0.6 at 600 nm. Then, bacteria were induced by 150 μM of IPTG and kept shaking at 28°C for four hours. Cells were harvested by centrifugation and resuspended in lysis buffer (sodium phosphate buffer pH-8 containing 1% Triton X-100 and 10 mM of dithiothreitol, lysozyme (1mg/ml)) and incubated on ice for 30 min. Next, the crude lysate was sonicated and centrifuged at 12000 rpm and 4°C for 10 min, and the supernatant containing the protein was collected for *in vitro* incubation with the substrate. Synthetic substrates were purchased from Buchem B. V. (Apeldoorn, Netherlands). Substrates were prepared according to Wang *et al*. (2019). The dried substrate was resuspended in an ethanolic detergent mixture of 0.4% (v / v) Triton X-100. The mixture was then dried using a vacuum centrifuge to produce an apocarotenoid-containing gel. The gel was resuspended in incubation buffer (2 mM tris 2-carboxyethylphosphine, 0.4 mM FeSO_4_ and 2 mg/ml catalase in 200 mM Hepes/ NaOH, pH 8). OsZAS2 crude cell lysate, 50 μl of the soluble fraction of overexpressing cells, was added to the assay. The assay was incubated for 4 h shaking at 140 rpm at 28°C in dark. The reaction was stopped by adding two volumes of acetone and the lipophilic compounds were separated by partition extraction with petroleum ether: diethyl ether 1: 4 (v /v), dried, and resuspended in methanol for HPLC analysis. *In vitro* product were run on an Ultimate 3000 UHPLC system with a YMC Carotenoid C30 column (150 × 3.0 mm, 5 μm) following the parameters described in Wang *et al*. (2019).

### Subcellular localization

The *35S::OsZAS2:mNeongreen* constructed by amplifying the coding sequence of *OsZAS2* using specific primers (Supplemental Table S1). The PCR product subcloned into the pDONR221 entry vector by BP recombination reaction. Then, *OsZAS2::pDONR221* were fused into pB7FWG2,0 by Gateway cloning. The construct was introduced into *Agrobacterium tumefaciens* strain GV3101 by electroporation. Tobacco (*Nicotiana benthamiana*) infiltration was performed as described by Aljedaani *et al*. (2021). *35S::OsZAS2-mNeongreen*, membrane protein marker (*35S::Lit6BmTurquoise2*) and p19 helper plasmid were co-infiltrated into the abaxial leave side of tobacco. The fluorescence expression was checked 3 days post infiltration by the confocal microscope. Leaf tissues of the infiltrated *N. benthamiana* was mounted with water on microscope slides and visualized by using a high-resolution laser confocal microscope (STELLARIS 8 FALCON, Leica). The signals of the mNeongreeen were detected at 488 nanometer (nm) excitation filter and mTurquoise2 excitation at 446 nm.

### Mycorrhization

Rice seeds of wild-type cv. Dongjin (DJ), *Oszas2* (*Oszas2-d* and *Oszas2-a*) were germinated in pots containing a sand and incubated for 10 days in a growth chamber under a 14 h light (23 °C)/10 h dark (21 °C). All genotypes were colonized with ~ 1000 sterile spores of *R. irregularis* DAOM 197198 (Agronutrition, Labège, France). Mycorrhizal plants were grown in sand and watered with a modified Long-Ashton (LA) solution containing 3.2 μM Na_2_HPO_4_·12H_2_O and were grown in a growth chamber as described before. Wild type and *Oszas2* mutant plants were sampled at 14 dpi and 50 dpi. To analyse *OsZAS2* gene expression profiles wild type plants were inoculated with a fungal inoculum *Funneliformis mosseae* (BEG 12, MycAgroLab, France) mixed (25%) with sterile quartz. Non-mycorrhizal and mycorrhizal plants were sampled at 7 and 21 dpi. For all the experiments, mycorrhizal roots were stained with cotton blue, and the level of mycorrhizal colonization was assessed according to Trouvelot *et al*. (1986) using MYCOCALC (http://www2.dijon.inra.fr/mychintec/Mycocalc-prg/download.html). For the molecular analyses, roots were immediately frozen in liquid nitrogen and stored at −80 °C.

### *In situ* hybridization and GUS staining

For the sample preparation and embedding, rice roots were fixed in 4% paraformaldehyde in PBS (phosphate buffered saline: 130 mM NaCl; 7 mM Na_2_HPO_4_; 3 mM NaH_2_PO_4_, pH 7.4) overnight at 4°C. To facilitate the fixation, the samples were placed under vacuum for the first 15 to 30 min. Then the tissue was dehydrated in successive steps, each of 30 to 60 min duration, in 30%, 50%, 70%, 80%, 95%, and 100% ethanol and 100% Neo-Clear™ (Xylene substitute, Sigma-Aldrich). Finally, the samples were embedded in paraffin wax (Paraplast plus; Sigma) at 60°C. Sections of 7 to 8 mm were then transferred to slides treated with 100 mg/mL poly-L-Lys (Sigma) and dried on a warm plate at 40 °C overnight. In parallel, DIG-labeled RNA probes were synthesized starting with 1 mg of PCR-obtained template (Langdale, 1993). DIG-labeled riboprobes (antisense and sense probes) were produced with DIG-UTP by *in vitro* transcription using the Sp6 and T7 promoters according to the manufacturer’s protocol (RNA-labeling kit; Roche). The sections were treated as follows: deparaffinized in Neo-Clear™, rehydrated through an ethanol series, treated with 0.2 M HCl for 20 min, washed in sterile water for 5 min, incubated in 2X SSC for 10 min, washed in sterile water for 5 min, incubated with proteinase K (1 mg/mL in 100 mM Tris-HCl, pH 8.0, 50 mM EDTA; Roche) at 37 °C for 30 min, washed briefly in PBS, and then treated with 0.2% Glycine in PBS for 5 min. After two rinses in PBS, slides were incubated in 4% paraformaldehyde in PBS for 20 min, washed in PBS (2, 3, 5 min), and then dehydrated in an ethanol series from 30% to 100%. Hybridizations were carried out overnight at 55 °C with denatured DIG-labeled RNA probes in 50% formamide, 20X SSC, 20% SDS, 50 mg/mL tRNA, 40 μg/ml Salmon Sperm DNA. Slides were then washed twice in 1X SSC, 0.1% SDS at room temperature, and rinsed with 0.2X SSC, 0.1% SDS at 55 °C (2 3 10 min). After rinsing with 2X SSC for 5 min at room temperature, the nonspecifically bound DIG-labeled probe was removed by incubating in 10 mg/mL RNase A in 2X SSC at 37 °C for 30 min. Slides were then rinsed twice in 2% SSC before proceeding to the next stage. The hybridized probe was detected using an alkaline phosphatase antibody conjugate (Roche). After rinsing in TBS (100 mM Tris-HCl, pH 7.5, 400 mM NaCl) for 5 min, slides were treated with 0.5% blocking reagent in TBS for 1 h, incubated for 2 h with the anti-DIG alkaline phosphatase conjugate diluted 1:500 in 0.5% BSA Fraction V in TBS, and then washed in TBS (3 x 5 min). Color development was carried out according to Torres et al. (1995). The color reaction was stopped by washing in distilled water, and the sections were then dehydrated through an ethanol series, deparaffinized in Neo-Clear^™^, and mounted in Neomount (Merck) (Balestrini et al., 1997).

The GUS assay was performed on roots of pOsZAS2:GUS-L11andL18 colonized by *F. mosseae* and sampled at 35 dpi. Rice mycorrhizal root segments were cut and placed in single wells of a Multiwell plate and covered with freshly prepared GUS buffer (0,1 M sodium phosphate buffer pH 7.0, 5 mM K_4_Fe(CN)_6_, 5 mM K_3_Fe(CN)_6_, 0.3% Triton X, 0.3% x-Glc). To improve buffer penetration into the root segments, these were placed under vacuum for 10 min. Finally, samples were incubated at 37°C for 16 hours in the dark, de-stained with 70% ethanol, and observed under an optical microscope (Nikon Eclipse E300).

## Results

### OsZAS2 represents a separate clade in the ZAS CCD-subfamily

To clarify the phylogenetic relationship of *OsZASs (OsZAS, OsZAS-L1, OsZAS-L2*, and *OsZAS-L3*) with other plant ZAS genes, we first constructed a phylogenetic tree, using ZAS sequences from selected monocot and dicot species (Supplemental Table S2). This analysis divided the ZAS proteins into five clades (I-V) (Figure 1A). Three OsZAS enzymes, including OsZAS (Os09g0321200), OsZAS-L1 (Os08g0369800), and OsZAS-L2 (Os08g0371608) grouped in clade I, while OsZAS-L3 (Os06g0162550) clustered in clade II. Based on this analysis we renamed OsZAS-L1, OsZAS-L2, and OsZAS-L3 to OsZAS1b, OsZAS1c, and OsZAS2, respectively (Figure 1A). The clades III, IV, and V contain only enzymes from dicot species, which we called ZAS3, ZAS4, and ZAS5.

**Figure 1.**
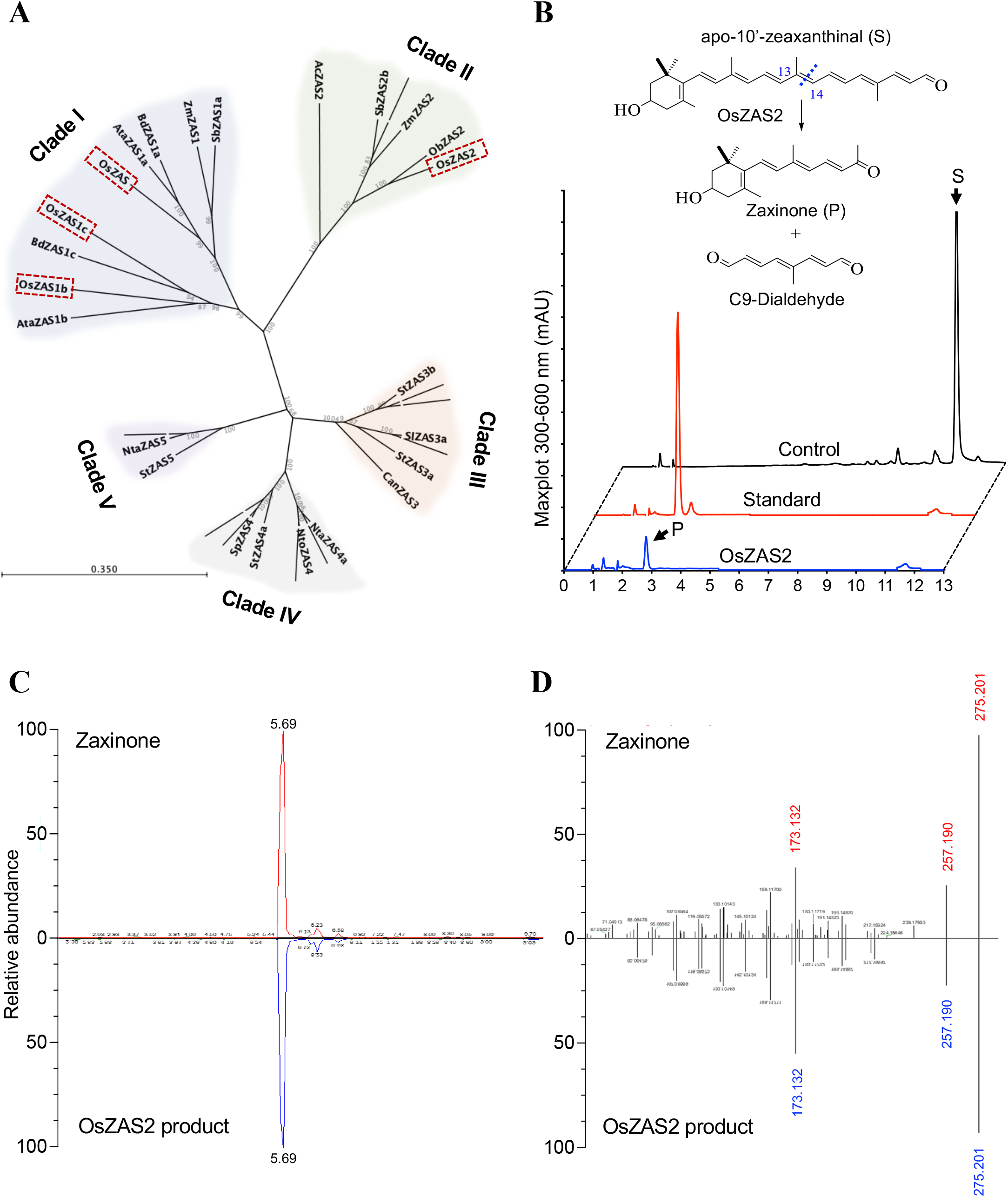
Phylogenetic analysis of ZAS enzymes and analysis of OsZAS2 enzymatic activity. A, Phylogenetic tree analysis of ZAS orthologues from selected monocot and dicot plants, showing bootstrap values on nodes of each cluster. Dashed rectangles represent rice ZAS members. B, HPLC chromatogram of *in vitro* incubation of OsZAS2 with apo-10’-zeaxanthinal (I) yielded zaxinone (II) and a presumed C_9_-dialdehyde (not shown). The maximum absorbance (mAU) peak for substrate and product is shown at 347 and 450 nm (mAU), respectively. C, Verification of OsZAS2 *in vitro* product, based on retention time, (D) accurate mass and MS/MS pattern and in comparison to zaxinone standard.

### OsZAS2 is a Zaxinone-Forming Enzyme

Next, we investigated the enzymatic activity of OsZAS2: we expressed OsZAS2 fused to maltose-binding protein (MBP) in *Escherichia coli* cells and incubated the soluble fraction of these cells with different apocarotenoids, i.e. β-apo-10’- (C_27_), 9-*cis*-β-apo-10’- (C_27_), β-apo-12’- (C_25_) and apo-8’-zeaxanthinal (3-OH-β-apo-8’-carotenal, C_30_) (Supplemental Figure S1). In addition, we incubated the MBP-OsZAS2 fusion with carotenoids, i.e. β-carotene, zeaxanthin, and lutein (Supplemental Figure S1). Finally, we performed *in vivo* activity test by expressing a thioredoxin-OsZAS2 fusion in β-carotene, zeaxanthin, and lycopene-accumulating *E. coli* cells. In all these assays, we only detected a conversion of apo-10’-zeaxanthinal (C_27_) that was cleaved by OsZAS2 at the C13, C14 double bond, yielding zaxinone (3-OH-β-apo-13-carotenone, C_18_) and a predicted C_9_-dialdehyde (Figure 1B). We confirmed the identity of zaxinone by UHPLC and LC-MS analysis, using a synthetic standard (Figure 1, C and D).

### OsZAS2 is a plastid enzyme expressed in roots and induced under low Pi

The ChloroP Server program (Emanuelsson et al., 1999) predicts the presence of a plastid transit peptide in the OsZAS2, indicating a plastid localization of this enzyme. To confirm this prediction, we transiently expressed *OsZAS2* cDNA fused with the sequence encoding mNeonGreen fluorescence protein under the control of the 35S promoter (*35S:OsZAS2:mNeonGreen*) in tobacco leaves epidermal cells, alone or together with the gene encoding the cell membrane specific Turquoise2 marker protein (*35S::Lit6BmTurquoise2*). As shown in Figure 2A, the green fluorescent signal of the OsZAS2 fusion overlapped with the red autofluorescence of chlorophyll A, demonstrating that OsZAS2 is a plastid enzyme.

**Figure 2.**
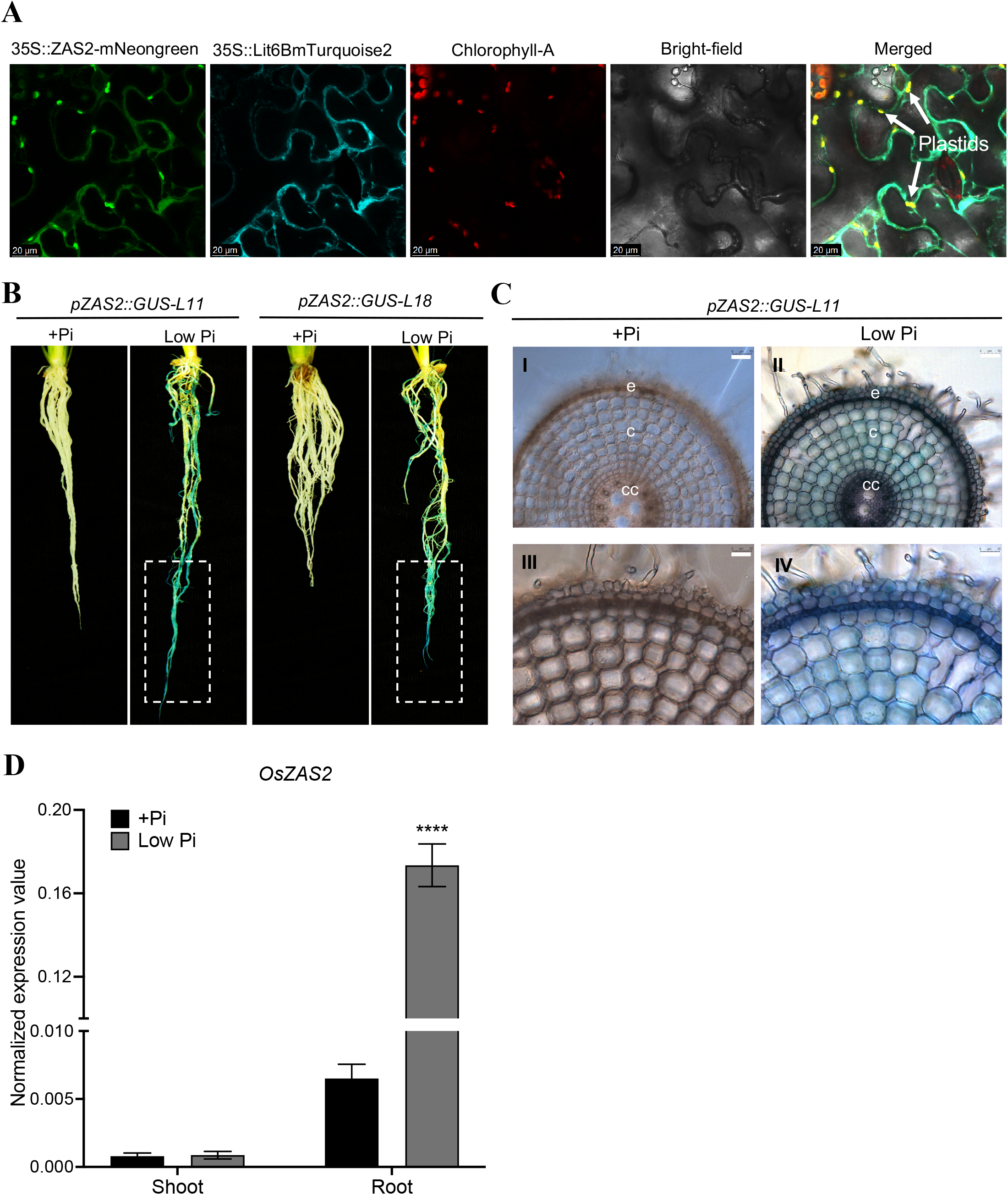
*OsZAS2* subcellular localization and expression pattern. A, Subcellular localization of OsZAS2 transiently expressed in tobacco epidermis leave tissue. cc, central cylinder; c, cortical cells; e, epidermal cells. B, GUS staining of roots of two independent *pZAS2:GUS* reporter lines (*pZAS2::GUS-L11, pZAS2::GUS-L18*) under normal (+Pi) and low Pi conditions. Dash rectangle emphasizes the root tip. C, Cross-section of *pZAS2:GUS11* line primary root (bars in pictures I, and II correspond to 25 μm, and bars in pictures III, and IV correspond to 50 μm). D, Normalized expression value of OsZAS2 under normal (+Pi) and low Pi conditions in root and shoot tissue of 21-day-old rice plants. Values in (D) are means ±SD (n = 4). Student’s t-test used for the statistical analysis (****P ≤ 0.0001).

To determine the expression pattern of *OsZAS2*, we generated the *GUS* reporter lines *pOsZAS2::GUS11* and *pZAS2::GUS18* by fusing a 1.2 kb upstream *OsZAS2* fragment to *GUS* and transforming the resulting *pZAS2-MCD162* plasmid into the rice. The staining of the two reporter lines showed that *OsZAS2* expression significantly increased under low Pi conditions, while the GUS signal was not detectable at all under normal conditions (Figure 2B). Moreover, the GUS signal increased significantly towards the tip of the primary and crown roots. The RT-qPCR analysis also showed that the *OsZAS2* transcript level increased about twenty-fold under low Pi compared to normal (+Pi) conditions in roots (Figure 2D). We did not detect the GUS signal in shoots, neither under nor under normal conditions (data not shown). We further investigated *OsZAS2* presence at the cellular level using cross-sectioning. Using a confocal microscope, we detected a strong GUS signal in the exodermis of primary roots, while the cortex, epidermis, and other root tissues showed only mild signals (Figure 2C).

### CRISPR/Cas9-generated *Oszas2* mutant lines show severe growth defects

Next, we generated *Oszas2* mutant knock-out lines in the Dongjin variety by employing CRISPR/Cas9. For this purpose, we used two guide RNAs, gRNA1 and gRNA2, that target coding sequence in exon 1 and exon 2, respectively (Figure 3A). We identified three independent *Oszas2* mutants, *zas2-d, zas2-g*, and *zas2-a*, with mutations causing premature stop codons in the first and second exon, respectively (Figure 3A). To validate the function of OsZAS2 as a zaxinone synthesizing enzyme *in planta*, we quantified zaxinone in roots of the three mutant lines grown hydroponically under normal Pi supply. Zaxinone content was reduced up to 45% in *Oszas2* mutant lines compared to wild-type (Figure 3C). Next, we grew the mutant lines in the soil and characterized them at the seedling and mature stages. *Oszas2* mutants displayed shorter roots and shoots and a striking reduction in root and shoot biomass (Figure 3, E-H). At the seedling stage, *Oszas2* lines produced one tiller on average, while the wild-type developed three (Figure 3D). The low-tillering and reduced shoot biomass phenotypes remained pronounced after growing the mutants for three months in GH and caused a significant reduction in grain weight per plant, compared to the corresponding wild type (Figure 4, A-E).

**Figure 3.**
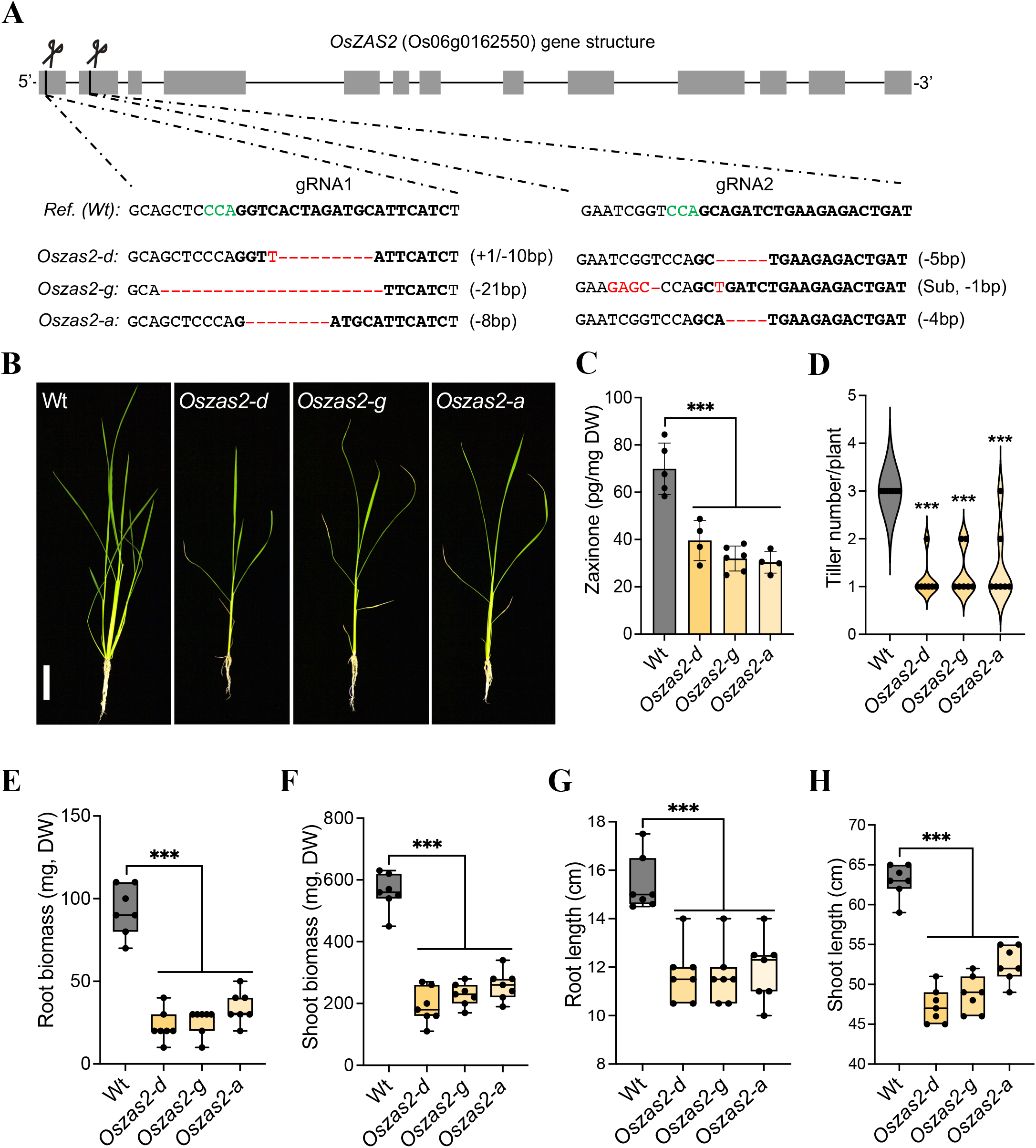
Characterization of CRISPR/Cas9-mediated *Oszas2* mutant lines at the seedling stages. A, Schematic representation of three individual mutations of *OsZAS2* gene generated by CRISPR-Cas9. B, The seedling phenotype of wild-type (Wt, Dongjin) and three independent *Oszas2* mutants. The scale bar in the picture represents 7.5 cm. C, Quantification of zaxinone content in Wt and *Oszas2* mutants roots. (D-H) Root biomass (D), shoot biomass (E), root length (F), shoot length (G), and tiller number (H) of the Wt and *Oszas2* mutants are shown in (B). Values in (C-H) are means ±SD (n ≥ 4). Student’s t-test used for the statistical analysis (***P ≤ 0.001).

**Figure 4.**
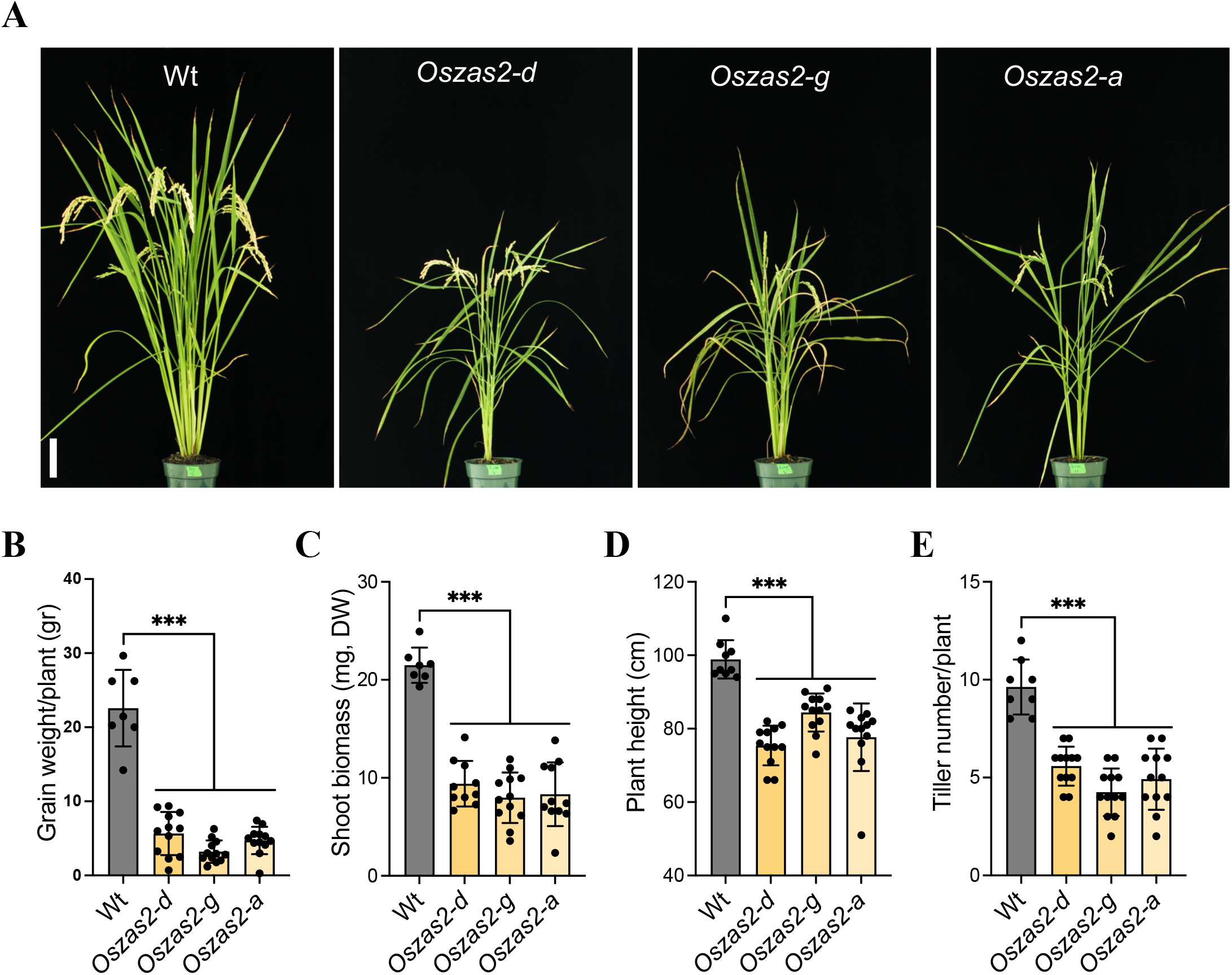
Characterization of *Oszas2* mutant lines at the maturing stage. A, The picture of the three-month-old Wt and *Oszas2* mutants grown in the greenhouse. The scale bar graph in the picture represents 7.5 cm. (B-E) Grain weight per plant (B), shoot biomass (C), plant height (D) and tiller number (E) of the Wt and *Oszas2* mutants represented in (A). Values in (b-e) are means ±SD (n ≥ 7). Student’s t-test was applied for the statistical analysis (***P ≤ 0.001).

### *OsZAS2* is involved in AM colonization

The expression analysis of *OsZAS2* on whole roots at early and late stages of AM colonization revealed an up-regulation at 21 dpi (days post-inoculation) (Supplemental Figure S2A), when arbuscules are present, as witnessed by the abundance of the AM-inducible plant marker *OsPT11* transcript (Paszkowski et al., 2002) (Supplemental Figure S2B). To obtain deeper insights into the spatial expression of *OsZAS2* during mycorrhization, we mycorrhized *pZAS2::GUS* reporter lines and monitored the GUS signal. Interestingly, we detected GUS activity only in arbusculated cells (Figure 5A; Supplemental Figure S2C), and did not observe any signal in any other root cells, including cortical cells and cells crossed by fungal hyphae. We further validated this observation by using *in situ* hybridization assays on mycorrhizal roots of wild-type plants. *OsZAS2* mRNA exclusively accumulated in cells with fully developed arbuscules, in which we detected a strong chromogenic signal (Figure 5B). We did not observe any signal in non-colonized cells or upon using the *OsZAS2* sense probe (Figure 5B).

**Figure 5.**
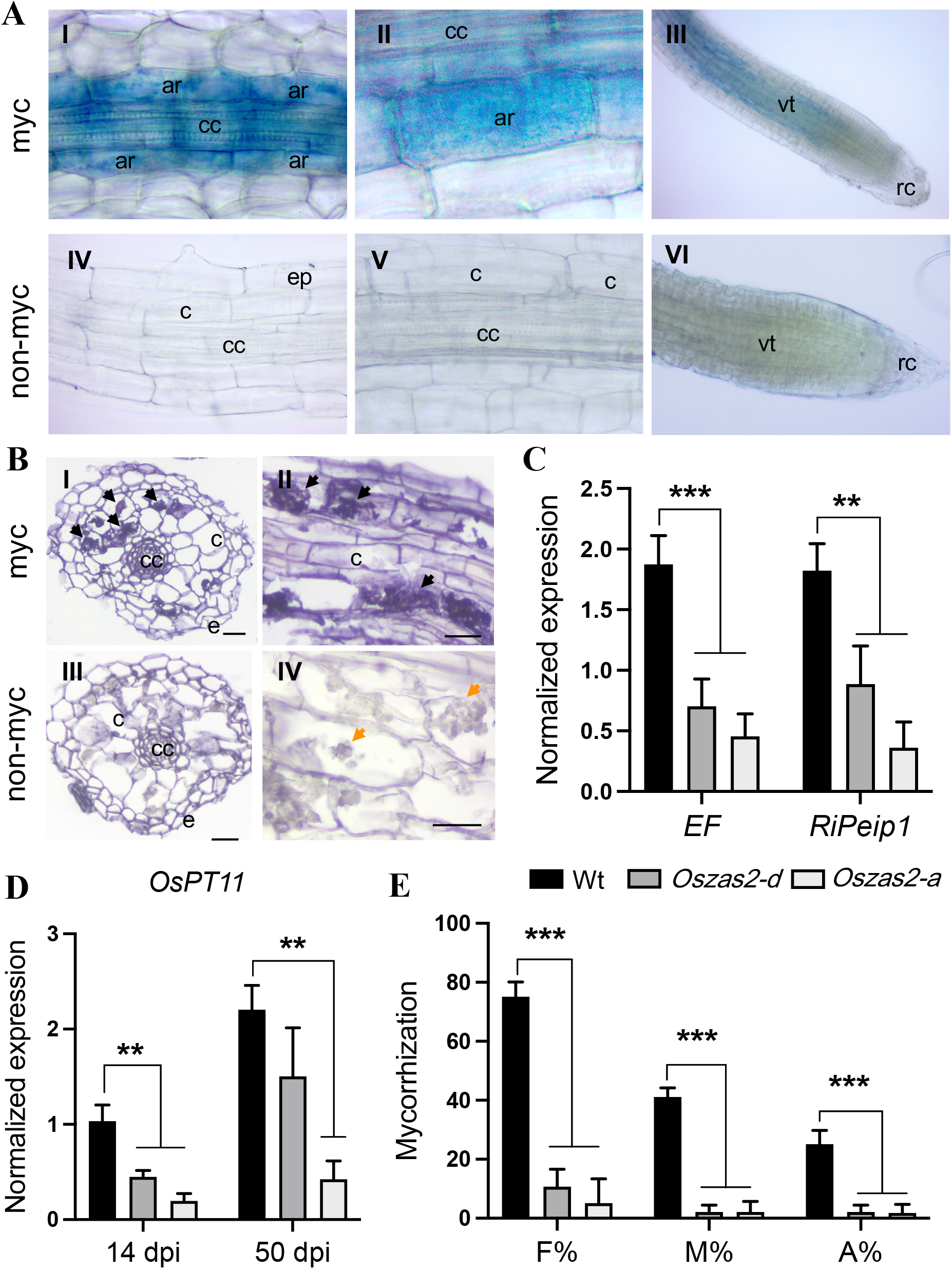
*OsZAS2* is required for arbuscular mycorrhizal (AM) establishment. A, GUS staining of roots of pZAS2:GUS-L18 reporter line inoculated (I, II, III) for 35 days with *F. mosseae* and non-inoculated (IV, V, VI). B, Localization of *OsZAS2* mRNA in sections from differentiated regions of inoculated roots by cold *in situ* hybridization. Section of mycorrhizal roots treated with *OsZAS2* antisense probe (I, II) and sense probe (III, IV). Black arrows represent a strong chromogenic signal, which mirrors the presence of the *OsZAS2* transcripts, is visible in arbuscule-containing cells (II). Orange arrows indicate the arbuscule-containing cells that are not labelled (IV). C, Relative expression of fungal genes; *RiEF* and *RiPeip1* in Wt and *Oszas2* mutants at 50 dpi. D, Relative expression of *OsPT11* at 14 and 40 dpi in Wt and *Oszas2* mutants. E, Frequency of mycorrhizal colonization (F%), the intensity of colonization (M%), and a total number of arbuscules (A%) in Wt and *Oszas2* mutants at 50 dpi (days post-inoculation). cc, central cylinder; c, non-colonized cortical cells; e, epidermal cells. Bars correspond to 50 μm. Values in (C-E) are means ±SD (n ≥ 3). Student’s t-test was applied for the statistical analysis (**P ≤ 0.01; ***P ≤ 0.001).

To determine the impact of *OsZAS2* on AM symbiosis, we mycorrhized the *Oszas2* mutant lines and assessed the colonization level by morphological analysis and by monitoring the transcript abundance of *OsPT11* at two-time points (14 dpi and 50 dpi) and of the fungal genes *RiEF* and *RiPeip1* (Fiorilli et al., 2016) at 50 dpi. Both mutant lines showed a lower frequency of mycorrhizal colonization (F%), intensity of colonization (M%), and total number of arbuscules (A%), compared to WT plants (Figure 5E). Molecular analysis confirmed these results: the expression level of fungal and plant genes was significantly lower in the mutant lines (Figure 5, C and D).

### SL biosynthesis increased in *Oszas2* mutants

The low-tillering phenotype of *Oszas2* mutants (Figure 3D; Figure 4E) indicated that they may have higher SL content. To test this hypothesis, we profiled their SLs in both roots and root exudates under low Pi conditions. We also analyzed the expression level of SL biosynthetic genes in roots. In roots, the contents of a non-canonical SL, a tentative 4-oxo-methylcarlactonate (4-oxo-MeCLA), and of the canonical SL 4-deoxyorobanchol (4DO) were significantly increased in *Oszas2* lines compared to wild-type (Figure 6A). As shown in Figure 6B, we also observed an increase in the transcript level of SL biosynthetic genes, i.e. *D27, CCD7, CCD8*, and *OsCO*. To confirm the increase in SLs, we conducted *a Striga* seed germination assay with root extracts. Extracts of *Oszas2* roots showed a significant germination rate (around 60%), compared to those of wild-type (around 42%, Figure 6D). In root exudates, both canonical (4DO, Orobanchol) and non-canonical (4-oxo-MeCLA) SLs were significantly increased in *Oszas2* mutants, compared to wild-type (Figure 6C). Here again, we performed a *Striga* seed germination assay, in which *Oszas2* mutant lines displayed about 15% higher activity compared to wild-type (Figure 6D), which confirms the LC-MS quantification.

**Figure 6.**
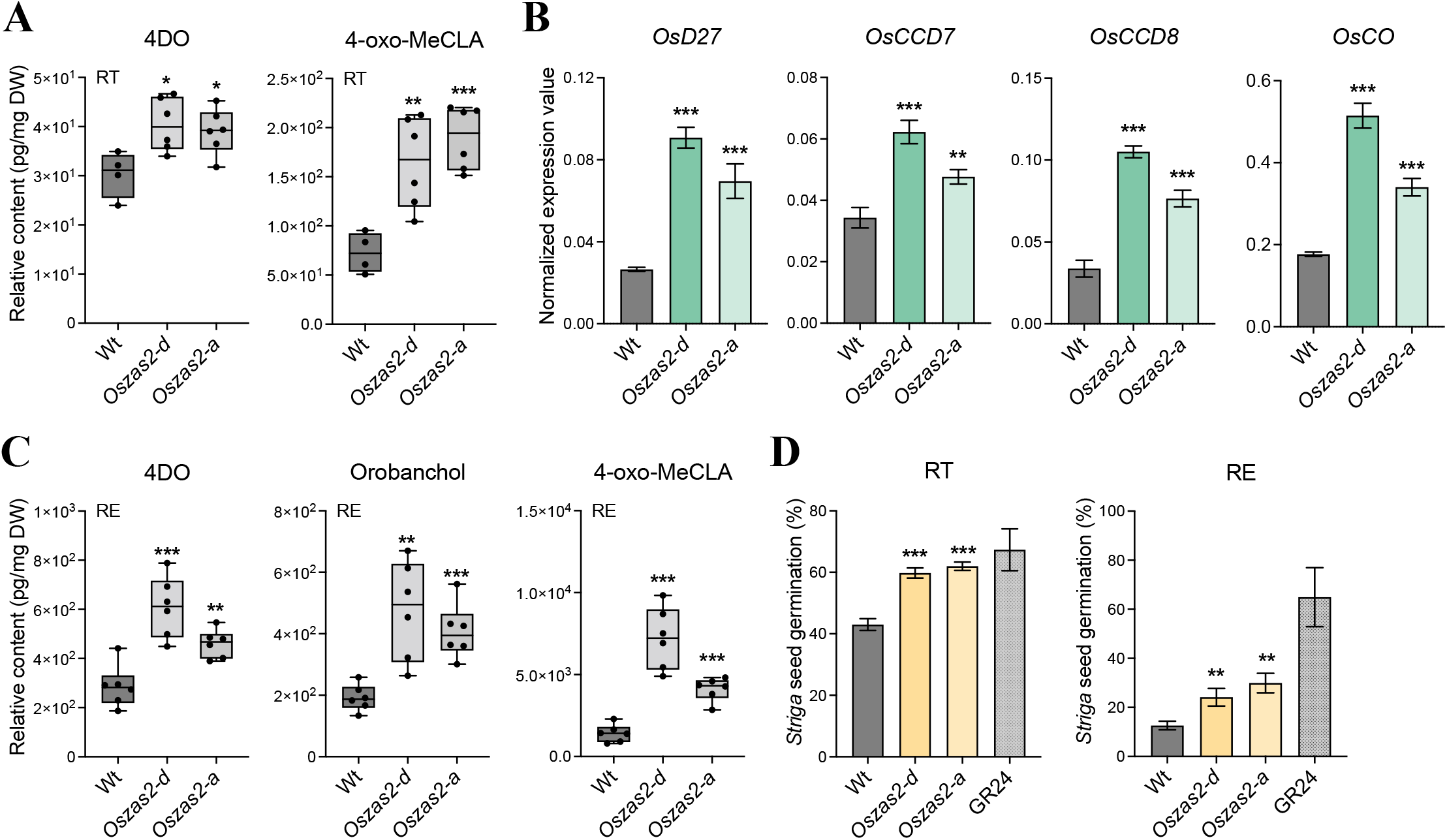
SL biosynthesis increased in *Oszas2* mutants. A, Relative quantification of 4DO and 4-oxo-MeCLA in root tissue (RT) of *Oszas2* mutants. B, Normalized expression value of SL biosynthetic genes in *Oszas2* mutants. C, Relative quantification of canonical (4DO, Orobanchol) and non-canonical (4-oxo-MeCLA) SL in root exudate (RE) of *Oszas2* mutants. D, *Striga* seed germination assay conducted with root exudate and tissue of *Oszas2* mutants. For both root exudate and tissue bioassay, 1 μM of GR24 was used as control, which showed about 65% and 67% of *Striga* seed germination, respectively. Values in (B-C) are means ±SD (n ≥ 4). Student’s t-test was applied for the statistical analysis (*P ≤ 0.05; **P ≤ 0.01; ***P ≤ 0.001).

### Exogenous zaxinone application repressed SL biosynthesis in *Oszas2* mutant

Next, we treated three weeks old, hydroponically grown *Oszas2* seedlings (grown one week in Hoagland agar and two weeks in low Pi) with 5 μM zaxinone for 6h. As shown in Figure 7A, zaxinone treatment repressed transcript levels of the SL biosynthetic genes *D27*, *CCD7, CCD8*, and *OsCO* in *Oszas2-d* back to the wild-type level. Furthermore, as shown before (Wang et al., 2019; Wang et al., 2020), exogenous zaxinone application decreased the transcript level of SL biosynthetic genes in wild-type as well (Figure 7A). Zaxinone application also decreased the content of the non-canonical SL 4-oxo-MeCLA in roots and root exudates of *Oszas2-d* and wild-type (Figure 7B). In addition, it caused a reduction in the level of the canonical SL orobanchol in root exudates of both *Oszas2-d* and wild-type (Figure 7C). Surprisingly, 4DO content was slightly increased upon zaxinone treatment in root tissues of both *Oszas2-d* and wild-type while it was not affected in root exudates (Supplemental Figure S3).

**Figure 7.**
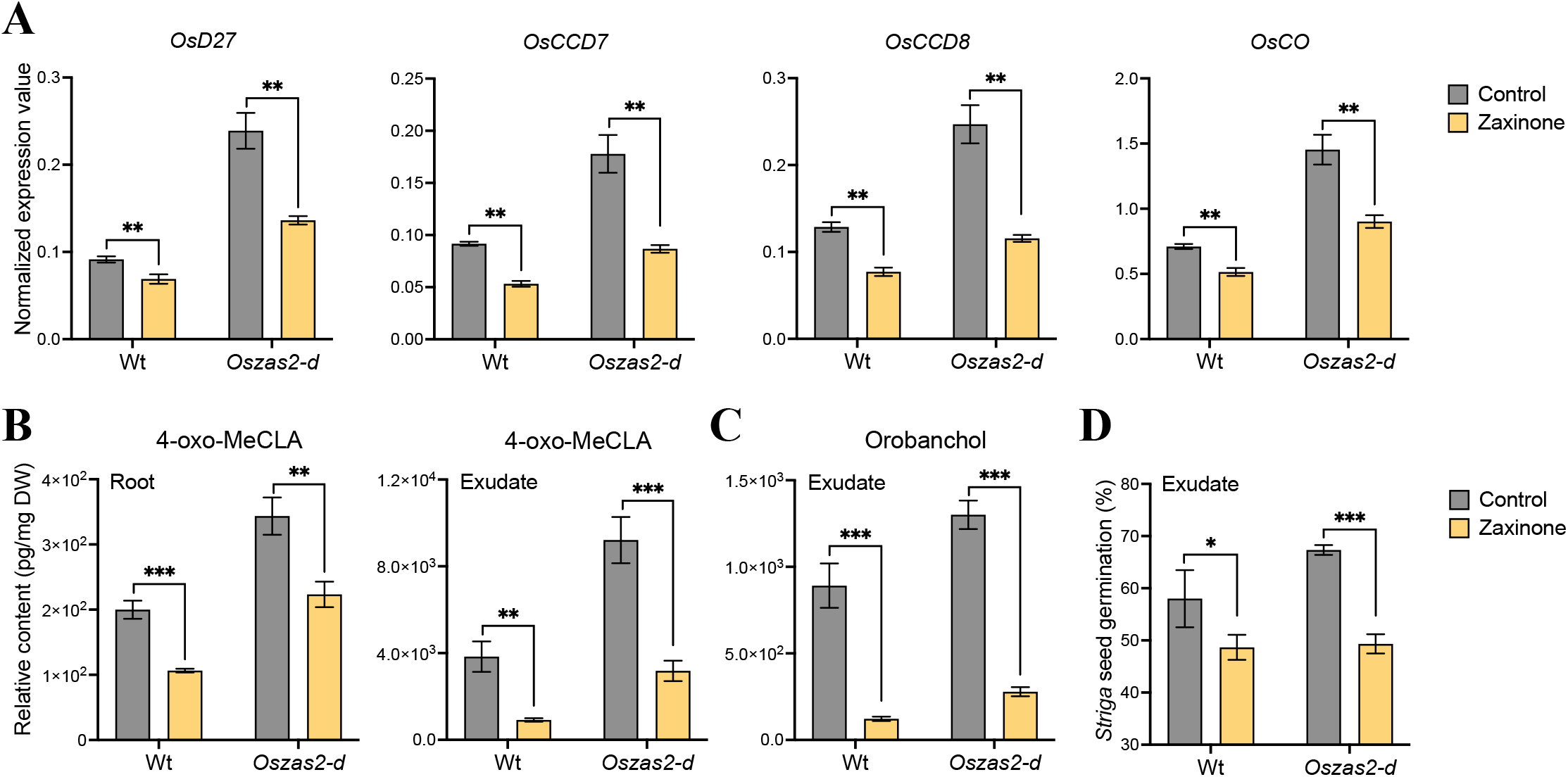
Zaxinone treatment reduced SL biosynthesis in *Oszas2* mutant. A, SL biosynthetic genes; *OsD27, OsCCD7, OsCCD8*, and *OsCO* expression in Wt and *Oszas2* mutant upon zaxinone (5 μM) treatment. B, Relative content of 4-oxo-MeCLA after zaxinone (5 μM) treatment in root tissue and exudate of Wt and *Oszas2* mutant. C, Relative content of Orobanchol after zaxinone (5 μM) treatment in root exudate of Wt and *Oszas2* mutant. D, *Striga* seed germination assay with exudate of Wt and *Oszas2* mutant upon zaxinone (5 μM) treatment. 1 μM of GR24 was used as control, which showed about 63% of *Striga* seed germination. Values in (A-D) are means ±SD (n ≥ 4). Student’s t-test was applied for the statistical analysis (*P ≤ 0.05; **P ≤ 0.01; ***P ≤ 0.001).

## Discussion

The identification of the first zaxinone synthase, the rice ZAS, unraveled the presence of a widely distributed plant CCD subfamily (Wang et al., 2019). Functional studies and characterization of a corresponding T-DNA insertion mutant demonstrated the importance of *ZAS* for plant growth, interaction with AM fungi, and SL homeostasis. Furthermore, exogenous treatment with zaxinone revealed the growth-promoting effect of this apocarotenoid and its impact on hormone homeostasis and sugar metabolism, indicating that it may be a candidate for a novel plant hormone (Wang et al., 2019; Wang et al., 2021). Rice contains three *OsZAS* homologs (Wang et al., 2019), called here *ZAS1b, ZAS1c*, and *ZAS2*, with unknown functions. The severe phenotypes of *zas* mutant indicate the importance of this gene and suggest that its activity cannot be compensated by that of its homolog(s), and that the latter may exert different function(s). Therefore, we were interested in investigating the biological function of these enzymes.

Phylogenetic analysis placed OsZAS2 in a clade different from that of ZAS, ZAS1b, and 1c, indicating a different biological role and maybe enzymatic activity (Figure 1A). Therefore, we focused in this study on OsZAS2. However, the enzymatic studies performed here demonstrate that OsZAS2 catalyzes the same reaction as ZAS, i.e. it converts apo-10’-zeaxanthinal into zaxinone (Figure 1B). Indeed, the enzyme did not convert other substrates tested, e.g. β-carotene, zeaxanthin, or different apocarotenoids, or produced other products, pointing to zaxinone-formation as its enzymatic function. The same activity was reported for OsZAS. However, OsZAS cleaved, in addition to apo-10’-zeaxanthinal, apo-12’- and apo-8’-zeaxanthinal, but with lower activity (Wang et al., 2019). This difference might be caused by a wider substrate specificity. However, it is also possible that the heterologously expressed OsZAS is generally more active than ZAS2.

To explore the biological function of OsZAS2 *in planta*, we generated *Oszas2* knock-out lines using the CRISPR/Cas9 technology. Disrupting *OsZAS2* led to around a 40% decrease in roots zaxinone content. This decrease supports the *in vitro* enzymatic activity of OsZAS2 and suggests that OsZAS2 is an enzyme, besides OsZAS, responsible for zaxinone biosynthesis in rice. *Oszas2* mutants still contained a significant amount of zaxinone in roots. This could be due to the activity of OsZAS, which might compensate for the zaxinone production in the rice root. Nevertheless, zaxinone is common at higher levels in green tissues than in roots and is also present in plant species, such as Arabidopsis, which lacks *ZAS* genes, indicating that it can be synthesized via the alternative route(s) independent of ZAS enzymes (Wang et al., 2019; Mi et al., 2019; Ablazov et al., 2020). However, the phenotypes observed with *Oszas* and *Oszas2* suggest the importance of these enzymes and indicate the zaxinone content in roots is crucial for normal growth and development. We are currently generating *Oszas*/*Oszas2* double mutants, which could give us a hint about the involvement of other route(s) in zaxinone biosynthesis.

Zaxinoneis a negative regulator of SL biosynthesis in rice (Wang et al., 2019). Indeed, the *Oszas* mutant contained and released higher amounts of SLs (Wang et al., 2019) under Pi starvation, and this increase could be suppressed by exogenous zaxinone application. Based on its zaxinone-forming activity and the low-tillering phenotype of the *Oszas2* mutants, we assumed that loss of OsZAS2 may also cause an increase in SL content. Therefore, we quantified the SL and zaxinone content in *Oszas2* mutants under low Pi condition. Indeed, SL biosynthetic genes; *OsD27, OsCCD7, OsCCD8*, and *OsCO* transcript levels were up-regulated in *Oszas2*, compared to wild-type (Figure 6A). In parallel, both canonical and non-canonical SLs were significantly increased in roots and root exudates of *Oszas2* mutants (Figure 6, A and C), as confirmed by LC-MS analysis and by using the *Striga* seed germination bioassay. Interestingly, zaxinone content was not changed in *Oszas2* mutants under low Pi conditions (Supplemental Figure S4A), compared to the wild-type, albeit an increase in SL biosynthesis that is assumed to be caused by a decrease in zaxinone content. Since OsZAS is still functional in the *Oszas2* mutant, we hypothesized that it might compensate for the loss of *Oszas2* activity. In fact, *OsZAS* expression was up-regulated, but not its paralogs (*OsZAS1b* and *OsZAS1c*) in *Oszas2* mutant under low Pi condition (Supplemental Figure S4B). It might be speculated that changes in zaxinone content in certain root cells are crucial for regulating SL biosynthesis and that the increase in OsZAS activity may lead to a generally higher zaxinone content but cannot replace OsZAS2 in cells expressing this enzyme. Clarifying this point requires precise localization of both enzymes under low Pi conditions.

*Oszas2* mutants showed severe reduction in root and shoot biomass (Figure 3, E and F) and developed fewer tillers, compared to wild-type (Figure 3D). The retarded growth of *Oszas2* mutants demonstrates that *OsZAS2* is necessary for normal rice growth and development. In general, the phenotypes of the *Oszas2* mutants, reduced zaxinone content, retarded growth, and reduced tiller number, are similar to that of the *Oszas* mutant under normal conditions (Wang et al., 2019). Hence, it can be concluded that rice requires both *ZAS* and *ZAS2* genes to keep the root zaxinone concentration at a certain level, as well as for normal rice growth and development under normal condition.

We checked the expression pattern of OsZAS2 at tissue and cellular level using qRT-PCR and promoter-GUS-reporter lines. Similar to *OsZAS* (Wang et al., 2019), *OsZAS2* is expressed in roots and induced upon Pi starvation. A robust up-regulation of both *OZAS* and *OsZAS2* in rice roots in response to Pi starvation indicates their involvement in the plant’s response to Pi deficiency. Interestingly, analysis of the GUS reporter lines (*pOsZAS2::GUS11* and *pZAS2::GUS18*) demonstrated that the *OsZAS2* expression level was higher in root tips (Figure 2A). The cross-sectioning of the primary roots of the *pOsZAS2::GUS11* further showed that *OsZAS2* is highly expressed in exoderms. Here again, it would be very interesting to monitor *OsZAS* expression patterns at the cellular level to get insights into the function of *OsZAS* and *OsZAS2* and understand why both of them are important for proper rice growth.

In a previous work, we demonstrated that *zas* mutant showed a lower AM colonization level, compared to wild-type plants (Wang et al., 2019). Moreover, we revealed that the *ZAS* gene family is absent in genomes of non-AM host species, such as Arabidopsis (Wang et al., 2019; Fiorilli et al., 2019), suggesting a strong link between *ZAS* and AM symbiosis. To investigate the role of OsZAS2 in the different steps of AM colonization, we monitored its expression level during a time course experiment in mycorrhizal and non-mycorrhizal roots. Contrarily to *OsZAS* which was up-regulated during both early and later stages, *OsZAS2* was only up-regulated at the maximum of arbuscules formation (21 dpi), suggesting an involvement in arbuscules development/formation. This assumption is supported by *in situ* hybridization and using the *pZAS2::GUS* reporter lines: indeed both assays revealed that *OsZAS2* expression is localized in arbusculated cells. To further clarify the *OsZAS2* involvement during the AM symbiosis, we assessed the *Oszas2* mutant lines (*Oszas2-d* and *Oszas2-a*) colonization level at the morphological level and using molecular analyses. Although *Oszas2* mutants displayed in non-mycorrhizal condition a higher level of SLs in roots and root exudates, they showed a severe reduction of AM colonization level; however, no defects in arbuscules morphology were detected. A similar phenotype was also observed in the *Oszas* mutant (Wang et al., 2019). All the above data demonstrate that OsZAS2 is required to reach a correct level of AM colonization.

We assume that the growth retardation of the *Oszas2* mutants is more likely due to decreased root zaxinone levels under normal conditions. Therefore, we supplied *Oszas2* seedlings, grown in soil, with exogenous zaxinone at a concentration of 10 μM for 2 weeks. We observed an increase in the tiller number of *Oszas2-d* mutant upon zaxinone treatment but of that of the wild-type. In contrast, the tiller number of wild-type was not changed upon zaxinone application (Figure 8, A and B). Moreover, zaxinone treatment significantly increased root and shoot biomass and length of *Oszas2-d* (Figure 8, C-F). This result highlights again the importance of appropriate zaxinone concentrations for regular growth and development of rice and supports the function of this apocarotenoid as a growth-promoting metabolite.

**Figure 8.**
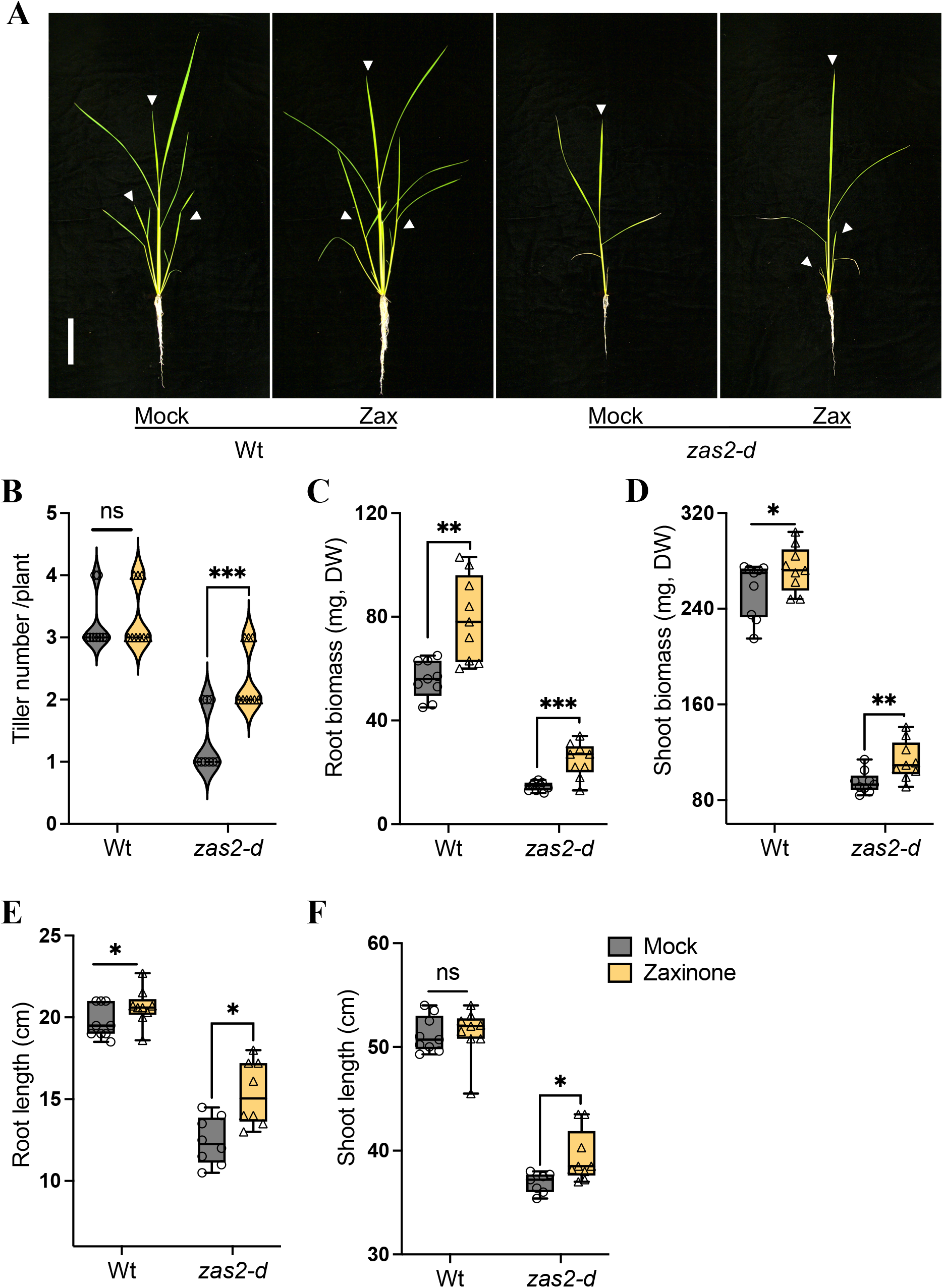
Exogenous zaxinone application rescued the growth defects of Oszas2 mutant. A, The images of the wild-type (Wt, Dongjin) and *Oszas2-d* mutant grown for two weeks in soil supplemented with 10 μM of zaxinone and tap water (0.01% acetone) as mock. The white bar represents 10 cm of scale. (B-F) Tiller number (B), root biomass (C), shoot biomass (D), root length (E), and shoot length (F) of the Wt and Oszas2 mutants are shown in (A). Values in (B-F) are means ±SD (n ≥ 7). Student’s t-test used for the statistical analysis (*P ≤ 0.05; **P ≤ 0.01; ***P ≤ 0.001; ns: not significant).

In conclusion, we revealed the function of *OsZAS2*, a new member of the *CCD* gene family, which is crucial for normal rice growth and development. Besides OsZAS, OsZAS2 contributes to zaxinone production in rice roots and is a further determinant of SL level. Moreover, it is involved in the regulation of mycorrhizal colonization levels. Thus, manipulation of OsZAS2 expression level could be a tool to modulate rice architecture and improve AM symbiosis.

## Acknowledgment

This work was supported by baseline funding and a Competitive Research Grant (CRG2017 and CRG 9) given to Salim Al-Babili from King Abdullah University of Science and Technology. We are thankful to Dr. Imran Haider for helping with rice transformation.

## Author Contributions

SA-B and AA conceived and designed the research; SA-B supervised the experiments. AA generated transgenic lines with the help of CHR. CV, VF, RB and LL planned and performed experiments concerning mycorrhization. MJ and AA conducted *Striga* seed germination assays. AB performed *in vitro* assays. AA characterized the transgenic lines with help of MJ. KXL and AA performed metabolite analysis with help of JYW. FA performed subcellular localization. IB helped with taking the microscope pictures. LB helped with plant growing and material preparation. AA analyzed the data and generated the figures and wrote the manuscript with the help of JYW, VF, CV, LL, and FA. SA-B edited and approved the article.

## Data Availability

The cDNA and promoter sequence of rice ZAS2 is available in NCBI under the accession number LOC107275952.

## Supporting information

**Supplemental Figure S1** Structures of carotenoids and apocarotenoids used as substrates in ZAS2 *in vitro* and *vivo* assays.

**Supplemental Figure S2** *OsZAS2* expression during arbuscular mycorrhizal (AM) establishment. A, *OsZAS2* expression pattern at a different stage of mycorrhizal colonization (*R. irregularis*). B, *OsPT11 (*AM marker gene) expression pattern at a different stage of mycorrhizal colonization (*R. irregularis*). C, GUS staining analysis of roots of *pZAS2:GUS-L11* reporter line inoculated (I, II, III) for 35 days with *F. mosseae* and non-inoculated (IV, V, VI). cc, central cylinder; c, non-colonized cortical cells; e, epidermal cells, dpi, day post-inoculation. Student’s t-test was applied for the statistical analysis (*P ≤ 0.05).

**Supplemental Figure S3** Relative content of 4DO after zaxinone (5 μM) treatment in root tissue and exudate of Wt and *Oszas2* mutant. Values in are means ±SD (n ≥ 5). Student’s t-test was applied for the statistical analysis (*P ≤ 0.05; **P ≤ 0.01; ns: not significant).

**Supplemental Figure S4** Zaxinone quantification and *OsZAS* genes expression analysis in *Oszas2* mutants under low Pi conditions. A, Quantification of zaxinone content in Wt and *Oszas2* mutants roots under low Pi condition. B, *OsZAS* genes expression analysis in Wt and *Oszas2-d* under low Pi condition. Values in (A-B) are means ±SD (n ≥ 3). Student’s t-test was applied for the statistical analysis (***P ≤ 0.001; ns, not significant).

**Supplemental Figure S5** gRNA targets of OsZAS2 were fused to tRNA sequences (Xie et al., 2015).

**Supplemental Table S1** Primer sequences used in these study.

**Supplemental Table S2** Distribution of ZAS members across monocot and dicot plants.

## References

Ablazov A, Mi J, Jamil M, Jia KP, Wang JY, Feng Q, Al-Babili S (2020) The apocarotenoid zaxinone is a positive regulator of strigolactone and abscisic acid biosynthesis in Arabidopsis roots. Frontiers in Plant Science. 578.

Ahrazem O, Gómez-Gómez L, Rodrigo MJ, Avalos J, Limón MC (2016) Carotenoid cleavage oxygenases from microbes and photosynthetic organisms: features and functions. International journal of molecular sciences, 17 (11):1781.

Aljedaani F, Rayapuram N, Blilou I (2021) A Semi-In Vivo Transcriptional Assay to Dissect Plant Defense Regulatory Modules. Modeling Transcriptional Regulation 203–214.

Akiyama K, Matsuzaki KI, Hayashi H (2005) Plant sesquiterpenes induce hyphal branching in arbuscular mycorrhizal fungi. Nature 435: 824–7.

Al-Babili S, Bouwmeester HJ (2015) Strigolactones, a novel carotenoid-derived plant hormone. Annual review of plant biology 66:161–86.

Alder A, Jamil M, Marzorati M, Bruno M, Vermathen M, Bigler P, Ghisla S, Bouwmeester H, Beyer P, Al-Babili S (2012) The path from β-carotene to carlactone, a strigolactone-like plant hormone. Science 335 (6074):1348–51.

Auldridge ME, Block A, Vogel JT, Dabney-Smith C, Mila I, Bouzayen M, Magallanes-Lundback M, DellaPenna D, McCarty DR, Klee HJ (2006) Characterization of three members of the Arabidopsis carotenoid cleavage dioxygenase family demonstrates the divergent roles of this multifunctional enzyme family. The Plant Journal 45 (6):982–93.

Auldridge ME, McCarty DR, Klee HJ (2006) Plant carotenoid cleavage oxygenases and their apocarotenoid products. Current opinion in plant biology 9 (3): 315–21.

Ballottari M, Alcocer MJ, D’Andrea C, Viola D, Ahn TK, Petrozza A, Polli D, Fleming GR, Cerullo G, Bassi R (2014) Regulation of photosystem I light harvesting by zeaxanthin. Proceedings of the National Academy of Sciences 111: E2431–E2438.

Bouvier F, Dogbo O, Camara B (2003) Biosynthesis of the food and cosmetic plant pigment bixin (annatto). Science 300 (5628): 2089–91.

Braguy J, Ramazanova M, Giancola S, Jamil M, Kountche BA, Zarban R, Felemban A, Wang JY, Lin PY, Haider I, Zurbriggen M (2021) SeedQuant: a deep learning-based tool for assessing stimulant and inhibitor activity on root parasitic seeds. Plant physiology, 186 (3): 1632–1644.

Bruno M, Beyer P, Al-Babili S (2015) The potato carotenoid cleavage dioxygenase 4 catalyzes a single cleavage of β-ionone ring-containing carotenes and non-epoxidated xanthophylls. Archives of biochemistry and biophysics. 572:126–33.

Bruno M, Hofmann M, Vermathen M, Alder A, Beyer P, Al-Babili S (2014) On the substrate-and stereospecificity of the plant carotenoid cleavage dioxygenase 7. FEBS letters 588 (9):1802–7.

Bruno M, Koschmieder J, Wuest F, Schaub P, Fehling-Kaschek M, Timmer J, Beyer P, Al-Babili S (2016) Enzymatic study on AtCCD4 and AtCCD7 and their potential to form acyclic regulatory metabolites. Journal of experimental botany 67 (21): 5993–6005.

Decker EL, Alder A, Hunn S, Ferguson J, Lehtonen MT, Scheler B, Kerres KL, Wiedemann G, Safavi-Rizi V, Nordzieke S (2017) Strigolactone biosynthesis is evolutionarily conserved, regulated by phosphate starvation and contributes to resistance against phytopathogenic fungi in a moss, Physcomitrella patens. New Phytologist 216 (2): 455–68.

Dhar MK, Mishra S, Bhat A, Chib S, Kaul S (2020) Plant carotenoid cleavage oxygenases: structure–function relationships and role in development and metabolism. Briefings in Functional Genomics 19 (1): 1–9.

Emanuelsson O, Nielsen H, Von Heijne G (1999) ChloroP, a neural network-based method for predicting chloroplast transit peptides and their cleavage sites. Protein Science 8 (5):978–84.

Felemban A, Braguy J, Zurbriggen MD, Al-Babili S (2019) Apocarotenoids involved in plant development and stress response. Frontiers in Plant Science 10: 1168.

Fiorilli V, Wang JY, Bonfante P, Lanfranco L, Al-Babili S (2019) Apocarotenoids: old and new mediators of the arbuscular mycorrhizal symbiosis. Frontiers in plant science 10: 1186.

Fraser PD, Bramley PM (2004) The biosynthesis and nutritional uses of carotenoids. Progress in lipid research 43 (3): 228–65.

Giuliano G, Al-Babili S, Von Lintig J (2003) Carotenoid oxygenases: cleave it or leave it. Trends in plant science 8 (4): 145–9.

Gomez-Roldan V, Fermas S, Brewer PB, Puech-Pagès V, Dun EA, Pillot JP, Letisse F, Matusova R, Danoun S, Portais JC (2008) Strigolactone inhibition of shoot branching. Nature 455 (7210): 189–94.

Gutjahr C, Parniske M (2013) Cell and developmental biology of arbuscular mycorrhiza symbiosis. Annual review of cell and developmental biology 29: 593–617.

Hashimoto H, Uragami C, Cogdell RJ (2016) Carotenoids and photosynthesis. Carotenoids in nature 111–39.

Hiei Y, Komari T (2008) Agrobacterium-mediated transformation of rice using immature embryos or calli induced from mature seed. Nature protocols 3 (5): 824–34.

Ilg A, Beyer P, Al-Babili S (2009) Characterization of the rice carotenoid cleavage dioxygenase 1 reveals a novel route for geranial biosynthesis. The FEBS journal 276 (3): 736–47.

Ilg A, Bruno M, Beyer P, Al-Babili S (2014) Tomato carotenoid cleavage dioxygenases 1A and 1B: Relaxed double bond specificity leads to a plenitude of dialdehydes, mono-apocarotenoids and isoprenoid volatiles. FEBS Open Bio 4: 584–93.

Ito S, Braguy J, Wang JY, Yoda A, Fiorilli V, Takahashi I, Jamil M, Felemban A, Miyazaki S, Mazzarella T (2022) Canonical Strigolactones Are Not the Tillering-Inhibitory Hormone but Rhizospheric Signals in Rice. BioRxiv

Chernys JT, Zeevaart JA (2000) Characterization of the 9-cis-epoxycarotenoid dioxygenase gene family and the regulation of abscisic acid biosynthesis in avocado. Plant physiology 124 (1): 343–54.

Jamil M, Kountche BA, Haider I, Wang JY, Aldossary F, Zarban RA, Jia KP, Yonli D, Shahul Hameed UF, Takahashi I (2019) Methylation at the C-3’ in D-ring of strigolactone analogs reduces biological activity in root parasitic plants and rice. Frontiers in plant science 10:353.

Jia KP, Dickinson AJ, Mi J, Cui G, Xiao TT, Kharbatia NM, Guo X, Sugiono E, Aranda M, Blilou I (2019) Anchorene is a carotenoid-derived regulatory metabolite required for anchor root formation in Arabidopsis. Science advances 5 (11): eaaw6787.

Jia KP, Baz L, Al-Babili S (2018) From carotenoids to strigolactones. Journal of experimental botany 69 (9): 2189–204.

Jia KP, Mi J, Ablazov A, Ali S, Yang Y, Balakrishna A, Berqdar L, Feng Q, Blilou I, Al-Babili S (2021) Iso-anchorene is an endogenous metabolite that inhibits primary root growth in Arabidopsis. The Plant Journal 107 (1): 54–66.

Liang MH, He YJ, Liu DM, Jiang JG (2021) Regulation of carotenoid degradation and production of apocarotenoids in natural and engineered organisms. Critical Reviews in Biotechnology 41 (4): 513–34.

Mi J, Jia KP, Wang JY, Al-Babili S (2018) A rapid LC-MS method for qualitative and quantitative profiling of plant apocarotenoids. Analytica chimica acta 1035: 87–95.

Moise AR, Al-Babili S, Wurtzel ET (2014) Mechanistic aspects of carotenoid biosynthesis. Chemical reviews 114: 164-93

Moreno JC, Mi J, Alagoz Y, Al-Babili S (2021) Plant apocarotenoids: from retrograde signaling to interspecific communication. The Plant Journal 105 (2): 351–375.

Nambara E, Marion-Poll A (2005) Abscisic acid biosynthesis and catabolism. Annu. Rev. Plant Biol. 56: 165–185.

Nisar N, Li L, Lu S, Khin NC, Pogson BJ (2015) Carotenoid metabolism in plants. Molecular plant 8 (1): 68–82.

Peleg Z, Blumwald E (2011) Hormone balance and abiotic stress tolerance in crop plants. Current opinion in plant biology 14 (3): 290–295.

Ramel F, Birtic S, Ginies C, Soubigou-Taconnat L, Triantaphylidès C, Havaux M (2012) Carotenoid oxidation products are stress signals that mediate gene responses to singlet oxygen in plants. Proceedings of the National Academy of Sciences 109 (14): 5535–40.

Rodriguez-Concepcion M, Avalos J, Bonet ML, Boronat A, Gomez-Gomez L, Hornero-Mendez D, Limon MC, Meléndez-Martínez AJ, Olmedilla-Alonso B (2018) A global perspective on carotenoids: Metabolism, biotechnology, and benefits for nutrition and health. Progress in lipid research. 70: 62–93.

Schwartz SH, Tan BC, Gage DA, Zeevaart JA, McCarty DR (1997) Specific oxidative cleavage of carotenoids by VP14 of maize. Science 276 (5320):1872–4.

Trouvelot A, Kough JL, Gianinazzi-Pearson V (1986) Estimation of vesicular arbuscular mycorrhizal infection levels. Research for methods having a functional significance. In Physiological and genetical aspects of mycorrhizae= Aspects physiologiques et genetiques des mycorhizes: proceedings of the 1st European Symposium on Mycorrhizae, Dijon, 1-5 July 1985. Paris: Institut national de le recherche agronomique, c1986.

Umehara M, Hanada A, Yoshida S, Akiyama K, Arite T, Takeda-Kamiya N, Magome H, Kamiya Y, Shirasu K (2008) Inhibition of shoot branching by new terpenoid plant hormones. Nature 455 (7210): 195–200.

Ha CV, Leyva-González MA, Osakabe Y, Tran UT, Nishiyama R, Watanabe Y, Tanaka M, Seki M, Yamaguchi S, Dong NV (2014) Positive regulatory role of strigolactone in plant responses to drought and salt stress. Proceedings of the National Academy of Sciences 111 (2): 851–6.

Vogel JT, Tan BC, McCarty DR, Klee HJ (2008) The carotenoid cleavage dioxygenase 1 enzyme has broad substrate specificity, cleaving multiple carotenoids at two different bond positions. J. Biol. Chem. 283: 11364–11373.

Wang JY, Alseekh S, Xiao T, Ablazov A, Perez de Souza L, Fiorilli V, Anggarani M, Lin PY (2021) Multi-omics approaches explain the growth-promoting effect of the apocarotenoid growth regulator zaxinone in rice. Communications biology 4 (1): 1–1.

Wang JY, Chen GT, Jamil M, Braguy J, Sioud S, Liew KX, Balakrishna A, Al-Babili S (2022) Protocol for characterizing strigolactones released by plant roots. STAR protocols 3 (2): 101352.

Wang JY, Haider I, Jamil M, Fiorilli V, Saito Y, Mi J, Baz L, Kountche BA, Jia KP, Guo X (2019) The apocarotenoid metabolite zaxinone regulates growth and strigolactone biosynthesis in rice. Nature communications. 10 (1): 1–9.

Wang JY, Jamil M, Lin PY, Ota T, Fiorilli V, Novero M, Zarban RA, Kountche BA (2020) Efficient mimics for elucidating zaxinone biology and promoting agricultural applications. Molecular plant 13 (11): 1654–61.

Wang JY, Lin PY, Al-Babili S (2021) On the biosynthesis and evolution of apocarotenoid plant growth regulators. In Seminars in cell and developmental biology 109: 3–11.

Wang W, Shi J, Xie Q, Jiang Y, Yu N, Wang E (2017) Nutrient exchange and regulation in arbuscular mycorrhizal symbiosis. Molecular plant 10 (9): 1147–58.

Waters MT, Gutjahr C, Bennett T, Nelson DC (2017) Strigolactone signaling and evolution. Annual review of plant biology 68: 291–322.

Xie K, Minkenberg B, Yang Y (2015) Boosting CRISPR/Cas9 multiplex editing capability with the endogenous tRNA-processing system. Proceedings of the National Academy of Sciences 112 (11): 3570–3575.

Xie X, Yoneyama K, Yoneyama K (2010) The strigolactone story. Annual review of phytopathology. 48: 93–117.

Zheng X, Giuliano G, Al-Babili S (2020) Carotenoid biofortification in crop plants: citius, altius, fortius. Biochimica et Biophysica Acta (BBA)-Molecular and Cell Biology of Lipids 1865 (11): 158664.

Zheng X, Yang Y, Al-Babili S (2021) Exploring the diversity and regulation of apocarotenoid metabolic pathways in plants. Frontiers in Plant Science 12.

